# A stimulus-contingent positive feedback loop enables IFN-β dose-dependent activation of pro-inflammatory genes

**DOI:** 10.1101/2022.08.11.503561

**Authors:** Catera L. Wilder, Diane Lefaudeux, Raisa Mathenge, Kensei Kishimoto, Alma Zuniga Munoz, Minh A. Nguyen, Aaron S. Meyer, Quen J. Cheng, Alexander Hoffmann

## Abstract

Type I interferons (IFN) induce powerful anti-viral and innate immune responses via the transcription factor, IFN-stimulated gene factor (ISGF3). However, in some pathological contexts type I IFNs are responsible for exacerbating inflammation. Here, we show that a high dose of IFN-β also activates an inflammatory gene expression program in contrast to IFN-λ3, a type III IFN, which elicits only the common anti-viral gene program. We show that the inflammatory gene program depends on a second, potentiated phase in ISGF3 activation. Iterating between mathematical modeling and experimental analysis we show that the ISGF3 activation network may engage a positive feedback loop with its subunits IRF9 and STAT2. This network motif mediates stimulus-specific ISGF3 dynamics that are dependent on ligand, dose, and duration of exposure, and when engaged activates the inflammatory gene expression program. Our results reveal a previously underappreciated dynamical control of the JAK-STAT/IRF signaling network that may produce distinct biological responses, and suggest that studies of type I IFN dysregulation, and in turn therapeutic remedies, may focus on feedback regulators within it.

**HIGHLIGHTS:**

- High dose IFN-β activates a pro-inflammatory gene program in epithelial cells.
- IFN-β, but not IFN-λ3, induces a second, potentiated phase in ISGF3 activity.
- ISGF3 induces its subunits to form a stimulus-contingent positive feedback loop.
- The positive feedback motif is required for the pro-inflammatory gene program.

## INTRODUCTION

Host immune defense mechanisms are coordinated by interferons (IFNs). IFNs elicit cell-intrinsic anti-viral activity as well as cell-extrinsic inflammatory responses leading to activation and recruitment of diverse immune cells [1-3]. The different families of IFNs, type I, II, and III, engage these defense mechanisms to different degrees. Type I IFNs are widely known for their anti-viral activity [4, 5], but in several pathological scenarios there is evidence for a role in triggering inflammation [6]. Type II IFN is less known for generating an anti-viral state, but for an enhanced activation potential state that allows macrophages to orchestrate recruitment of diverse effector cells and initiate adaptive immune responses [7]. Like type I, type III IFNs have a primary role inducing anti-viral activity, but there is less evidence for activating inflammatory mediators [8-10].

Both type I and type III IFNs activate the same JAK-STAT/IRF signaling pathway and transcription factor, ISGF3 [11, 12]. While type I IFNs have a broad range of effectiveness due to the ubiquitous expression of the IFNAR subunits on most cell types [13], type III IFNs are associated with specific tissues due to the cell type-specific expression of these cytokines and the cognate IFNλR1 receptor subunit [13]. The human type I IFN family comprises 13 subtypes of IFN-α, IFN-β, IFN-ε, IFN-κ, and IFN-ω. Type III IFNs comprise four members, namely IFN-λ1, -λ2, -λ3, and -λ4. Altogether, there are 21 IFNs that activate the same transcription factor. This remarkable diversity of ligands activating the same transcription factor raises the question of whether their biological roles are overlapping or distinct. Studies have shown that IFN-α subtypes share the capacity to elicit anti-viral states but differ in their cytostatic effects in a manner that correlates with their affinity for the IFNAR receptor [14, 15]. Among the type I IFNs, IFN-β has been shown to be particularly potent due to its high binding affinity and sustained life-span of the IFN-IFNAR complex [14, 16]. Among the members of type III IFNs, IFN-λ3 has the most potent anti-viral activity [17, 18].

While the roles of type I and III IFNs in cell intrinsic immunity have been well-defined, their regulation of cell extrinsic immune functions has been less well characterized. A link between type I IFNs and inflammation has been described in various infections and may involve immune cell recruitment via the expression of monocyte recruiting chemokines [6, 19-21]. For type III IFNs the evidence for initiating cell extrinsic inflammation functions is less clear [22-25]. In clinical settings, type I IFNs have been used to treat hepatitis B and COVID-19, but its adoption as an effective therapeutic has been hampered by its adverse side effects (e.g. flu-like symptoms, fatigue) associated with inflammatory cytokines [26-28]. In contrast, type III IFN treatments have limited adverse effects, comparable to that of placebo groups [28-30].

The differential propensity of type I and III IFNs to induce inflammation could be due to cell type-specific expression of cognate receptors, or differences in signaling responses. In neutrophils, which respond to both type I and III IFNs, a type I IFN-specific inflammatory gene expression response was reported but not characterized mechanistically [22]. However, for lung epithelial cells, key to initiating immune responses during respiratory infections, no such information is available. Importantly, both type I and III IFNs play a role in their cellular responses, as an influenza A viral challenge in primary mouse tracheal epithelial cells deficient in either IFNAR or IFNλR1 resulted in similar gene expression programs [31]. Hence, there is a need to investigate if and how IFN-type I and III induce distinct responses within the same cell type.

The molecular mechanisms of signal transduction are well understood. Upon binding their cognate receptors, type I and III IFNs activate the same kinases JAK1 and TYK2, which activate the same transcription factor, ISGF3. ISGF3 is responsible for inducing the expression of hundreds of IFN-stimulated genes (ISGs) [2, 11, 32]. ISGF3 is a trimer composed of the active, phosphorylated forms of STAT1 and STAT2 along with IRF9. The active trimer translocates to the nucleus where it binds to the promoters of ISGs to induce gene expression. While ISGF3 is the primary transcription factor induced by type I and III IFNs, others such as GAF, MAPK/ERK, and NFκB have also been implicated as IFN-inducible [33-36]. A mathematical model of the IFN signaling pathway has been established and used to investigate how signaling is affected by prior IFN exposure or conditioning [37, 38]. However, whether and how ISGF3 signaling is distinct in response to type I and III IFN stimulation has not been quantitatively examined.

Here, we investigated whether type I IFN-β and III IFN-λ3 generate distinct gene expression responses in lung epithelial cells. We identified a type I IFN-specific gene expression program that is characterized by inflammatory response mediators. Iterating between quantitative experimentation and mathematical modeling, we investigated the mechanism and report that the dynamic control of a single transcription factor, ISGF3, allows for the distinction. A secondary, potentiated phase of ISGF3 is responsible for the inflammatory gene expression program. We show that the potentiated ISGF3 activation phase is enabled by a stimulus-contingent positive feedback loop that is engaged at high doses and sustained durations of type I IFN-β. Our results indicate that stimulus-specific dynamic control of ISGF3 elicits distinct immune gene expression responses, revealing the importance of transcription network feedback control in the JAK-STAT/IRF signaling system for healthy immunity.

## RESULTS

### IFN-β elicits both anti-viral and pro-inflammatory gene expression programs

In order to determine if IFN-β and IFN-λ3 induce differential expression of ISGs, we produced replicate RNA-seq datasets from an extensive time course. MLE-12 lung epithelial cells were stimulated with either 10 U/ml IFN-β or 100 ng/ml IFN-λ3. Concentrations were selected such that the initial activation of the IFN signaling pathway was comparable as measured by ISGF3 activity at 30 min (Supplemental Fig. 1). RNA was isolated at 11 time points of stimulation (0, 0.5, 1, 2, 3, 4, 6, 8, 12, 24, 36 hours) and subjected to polyA+ mRNA sequencing (RNA-seq) for global transcriptome analysis. 345 genes were identified as inducible (FDR corrected p-value < 0.05 for IFN-β dataset). Principal component analysis (PCA) of the selected genes yielded two components that accounted for almost 90% of the variance (Fig. 1A). While the transcriptomes of IFN-β and IFN-λ3 time courses remain similar at early time points (0.5 and 1 hours), they diverge on the PCA plot at two hours and are well-separated at four hours. The number of induced genes was similar at the early and late time points, but at four hours, IFN-β showed 2.3 times as many induced genes as IFN-λ3 (FDR corrected p-value < 0.05 and log_2_(FC) ≥ 1) (Fig. 1B). Interestingly, even for genes induced by both stimuli at four hours, there was a higher magnitude of induction with IFN-β (Fig. 1C). However, by 24 hours many genes showed a slightly higher induction with IFN-λ3.

**Figure 1.**
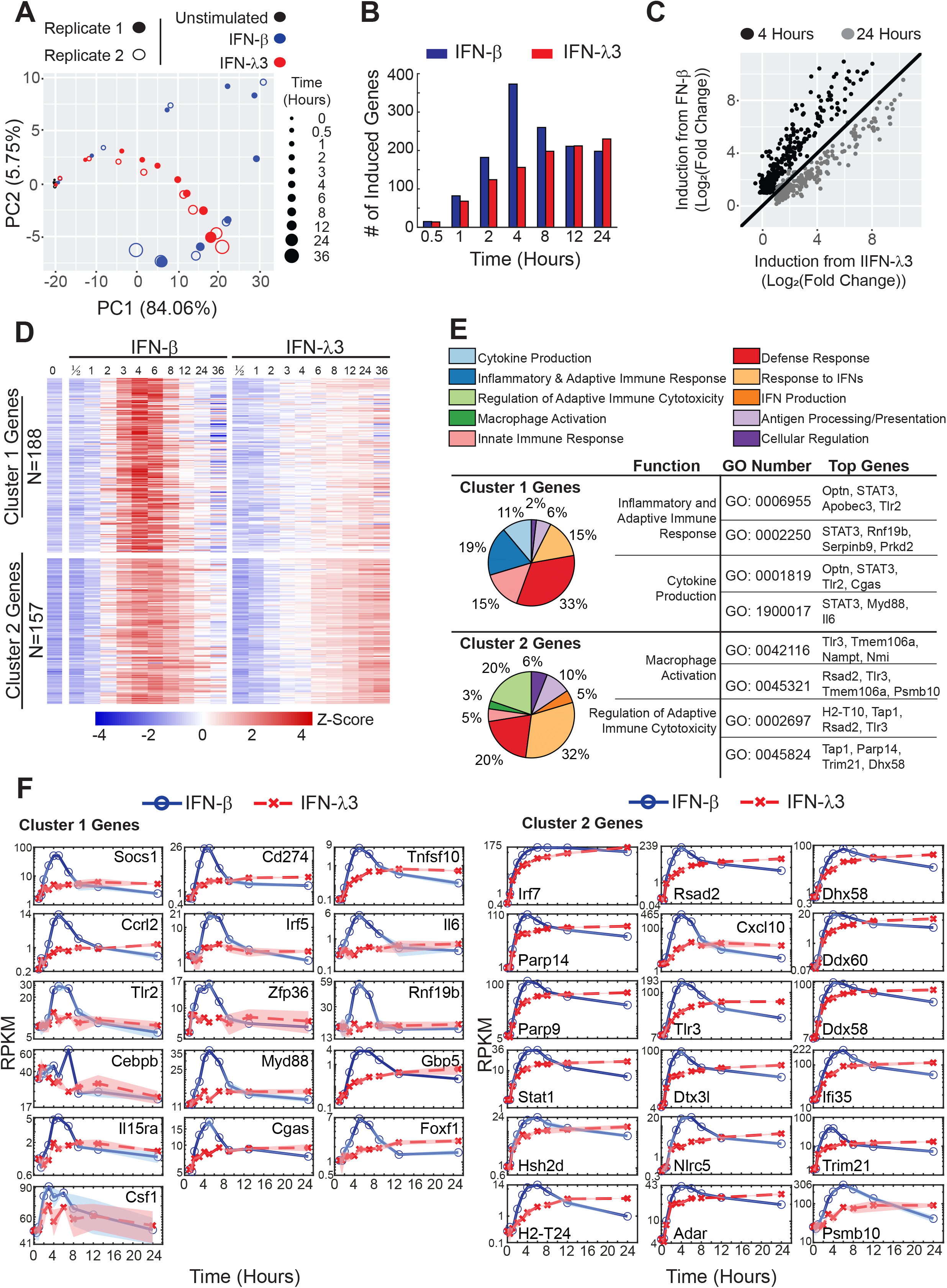
IFN-β, but not IFN-λ3, induces an inflammatory gene expression program. A. Principal component analysis (PCA) of interferon-response transcriptomes in MLE-12 cells. Replicate data sets (open and closed circles) for IFN-β (10 U/ml, blue) and IFN-λ3 (100 ng/ml, red) describe distinct trajectories over the two dimensional map of PC1 and PC2, at the indicated times of the time course (increasing circle size). Data are from two independent experiments. B. Bar graphs of the number of genes induced (log_2_(FC) ≥ 1, FDR corrected p-value ≤ 0.05) by IFN-β (blue) or IFN-λ3 (red) at each indicated time point. C. Scatter plot of fold induction in gene expression in response to IFN-λ3 versus IFN-β at an early time point (4 hours, black) and a late time point (24 hours, grey). D. Heatmap depicting the scaled mRNA expression time course of individual genes. The gene expression was averaged from two replicates and scaled (Z-score) for each gene in response to IFN-β and IFN-λ3 over time. Mean expression is depicted as white and values above or below the mean depicted as red or blue, respectively. Genes were clustered based on differential gene expression analysis. Genes in cluster 1 are differentially induced during at least one time point by IFN-β (FDR corrected p-value ≤ 0.05), but not IFN-λ3. Cluster 2 contains other genes that are induced by both IFN-β and IFN-λ3 during at least one time point compared to unstimulated. E. Pie charts of the biological functions of cluster 1 and 2 genes, as determined by Gene Ontology (GO) analysis. Biological programs are depicted in the indicated color code. Example biological functions that are specific to cluster 1 or 2 genes are listed with example GO ID numbers and top, over-represented genes from either cluster that mapped to the indicated GO ID. F. Line graphs depicting the mRNA expression time course (averaged from two replicates) induced by IFN-β (blue) or IFN-λ3 (red) of indicated genes from cluster 1 and 2. Genes contributing the highest variance to PC1 or PC2 (i.e. upper quartile) from the PCA analysis and that have important biological functions specific to cluster 1 or 2 from the GO analysis are shown plotted with connecting lines.

In order to assess the temporal dynamics at a single gene level, a heatmap was generated with each row representing expression of an individual gene (Fig. 1D). The selected 345 genes were grouped into two clusters based on induction by only IFN-β (cluster 1; FDR corrected p-value ≤ 0.05) or by both IFN-β and IFN-λ3 (cluster 2; FDR corrected p-value ≤ 0.05). Cluster 1 had 188 genes that generally showed transient activation with a peak at around four hours of IFN-β stimulation. The 157 genes in cluster 2 also showed transient induction in response to IFN-β but were also induced in response to IFN-λ3 with gene expression peaking at later time points (12, 24, 36 hours). Gene Ontology (GO) analysis identified potential cluster-specific biological functions such as “inflammatory and adaptive immune response” as well as “cytokine production” for cluster 1 genes and “regulation of adaptive immune cytotoxicity” as well as “macrophage activation” for cluster 2 genes (Fig. 1E).

To inspect the expression time course of specific genes with important cluster-specific biological functions in more detail, we selected genes based on top quartile loadings of principal component 2 (Fig. 1A), presence in cluster 1 (Fig. 1D), and association with cluster 1-specific GO functions (i.e. inflammatory & adaptive immune response and cytokine production). Line graphs of these cluster 1 genes confirmed the IFN-β-specific induction around the four-hour time point (Fig. 1F). A similar analysis was conducted for the top quartile genes contributing to variance in PC1 that are also found in cluster 2 and contribute to cluster 2-specific biological functions (i.e. regulation of adaptive immune cytotoxicity and macrophage activation). While many of these genes showed higher expression levels with IFN-β at four hours, with IFN-λ3 they reached almost equivalent expression levels at late time points.

### Differential ISGF3 temporal dynamics control gene expression programs

Since the temporal expression trajectories of cluster 2 genes were distinct when stimulated with either IFN-β or IFN-λ3, we hypothesized that IFN-specific temporal dynamics of the transcription factor ISGF3 was responsible. To examine this question, we measured ISGF3 activity in a time course of MLE-12 lung epithelial cells incubated with 10 U/ml IFN-β or 100 ng/ml IFN-λ3 using an electrophoretic mobility shift assay (EMSA) with an ISRE oligo probe using constitutive transcription factor NFY as control (Fig. 2A). We found three phases of ISGF3 activity during the IFN-β stimulation: an activation phase (0 to 60 minutes), a potentiated phase (60 to 240 minutes), and an inactivation phase (240 to 720 minutes). IFN-λ3-stimulated ISGF3 shared the activation phase, but lacked the potentiated phase, instead showing a trough (40 to 120 minutes), and re-activation phase (120 to 720 minutes). The IFN-β-specific potentiated phase is characterized by a more than a 2-fold increase of activity that peaks at four hours of stimulation.

**Figure 2.**
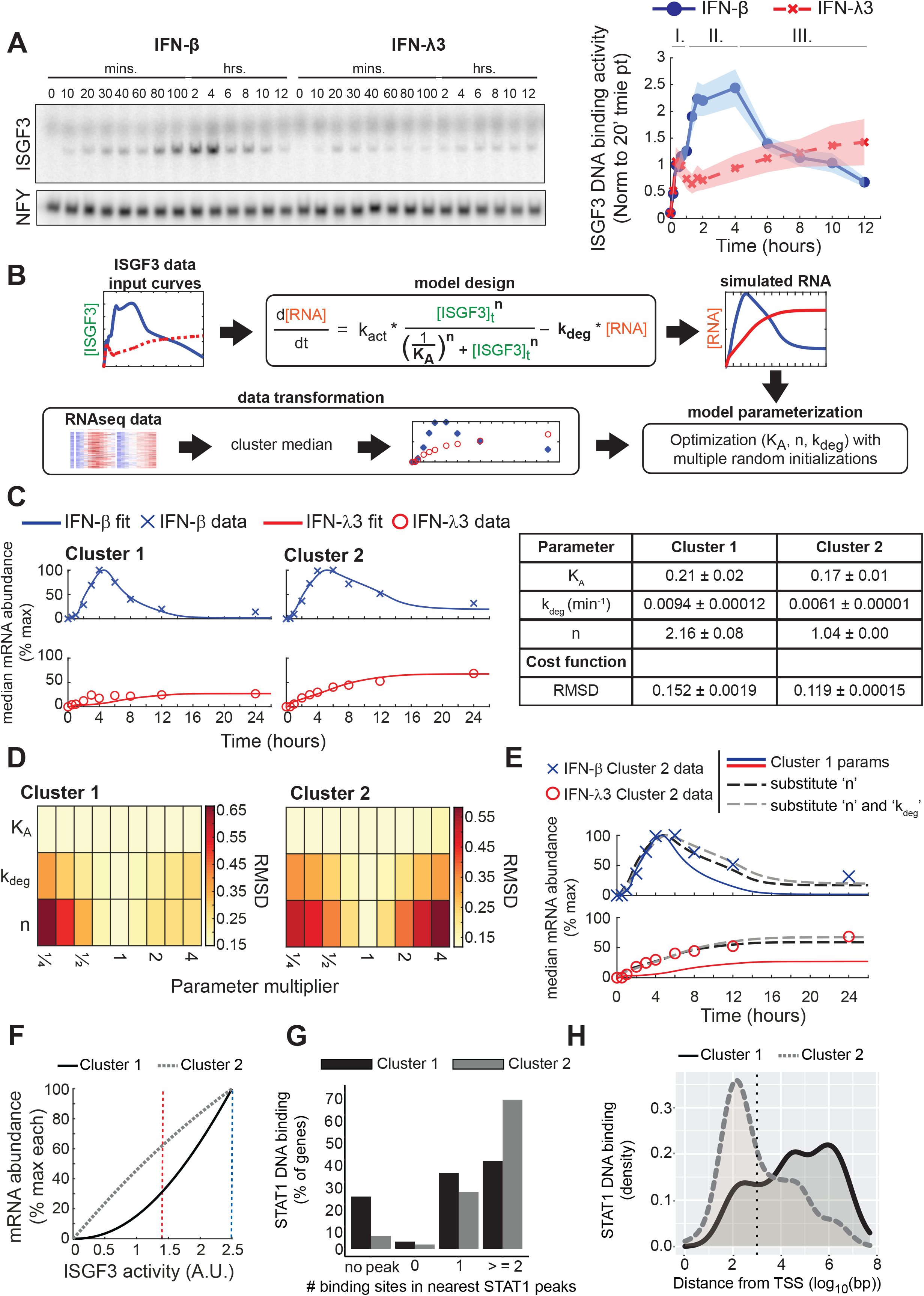
Stimulus-specific temporal dynamics of ISGF3 regulate ISG induction. A. ISGF3 activity revealed by an electrophoretic mobility shift assay (EMSA) using nuclear extracts prepared from MLE cells at indicated time points of IFN-β (10 U/ml, blue) and IFN-λ3 (100 ng/ml, red) stimulation. The activity was quantified and normalized to the ISGF3 activity at 20 minutes of IFN-β stimulation. Line graphs were plotted using connecting lines. Shaded areas on the line graphs depict the standard error of the mean (SEM) from at least five independent experiments. The three phases of ISGF3 activity are indicated above the graphs. B. Schematic of the ordinary differential equation (ODE) model and optimization pipeline. Quantified ISGF3 curves were used as input data (green text in equation) for an ODE-based model of mRNA expression with a Hill-based production term and degradation term. The model was parameterized by fitting the K_A,_ k_deg_, and n parameters (bold text in equation) to the median mRNA expression levels of cluster 1 and 2 time courses. C. Line graphs depicting the simulated best fit model (solid line) for genes in both clusters and the experimental median IFN-β (cross) and IFN-λ3 (open circle) induced-gene expression levels at each time point. Simulated and experimental data were normalized to the percentage of the maximum response of both stimuli. The optimized parameter sets and associated standard error of the mean of five optimization runs each represented by the best 10 of 50 resulting from random initializations for cluster 1 and cluster 2 are listed in the table. D. Heatmap of the root mean square deviation (RMSD) of simulated and experimental mRNA levels when changing parameters (K_A,_ k_deg_, and n) by the indicated value with the best fit models for cluster 1 and cluster 2. Good fits with lower RMSD values are depicted as a pale yellow while larger values indicating poor fits are dark red. E. Line graphs depicting the simulated best-fit model for cluster 1 (solid line) and the cluster 2 experimental median IFN-β- (cross) and IFN-λ3- (open circle) induced mRNA abundances at each time point. Simulations of the cluster 1 model after substituting the values of the Hill coefficient (black dotted line) or both Hill coefficient and the degradation rate constant (gray dotted line) with the value from the cluster 2 model. F. Line graphs of an ISGF3 dose-response curve for the cluster 1 and cluster 2 models. The mRNA level for different amounts of ISGF3 activity is plotted for the cluster 1 model (solid black line) and cluster 2 model (dotted gray line). The maximum amount of ISGF3 activity induced by either IFN-β (vertical dashed blue line) or IFN-λ3 (vertical dashed red line) are indicated in the graph. G. Bar graphs of the number of STAT1 binding motifs for genes in cluster 1 and 2. ChIP-seq analysis of STAT1 binding motifs were identified under STAT1 peaks. Cluster 1 and cluster 2 genes were mapped to the nearest STAT1 binding sites and the number of motifs under the nearest peak was counted and plotted. H. Density plots depicting the distance of genes in cluster 1 and 2 to the nearest STAT1 binding site. ChIP-seq analysis of the proportion of STAT1 peaks plotted based on their distance from the nearest gene. STAT1 binding less than or greater than 1,000 base pairs (vertical dotted black line) were classified as either proximal promoter or distal enhancer binding.

A simple model, composed of an ordinary differential equation (ODE), was used to determine if the temporal dynamics of ISGF3 activity might account for the measured cluster-specific expression dynamics in response to each stimulus. The ODE model simulates the change in mRNA abundance as a function of mRNA synthesis activated by the ISGF3 transcription factor with an activation constant (K_A_), Hill coefficient (n), and an mRNA degradation rate constant (k_deg_). The only time-dependent model variable is the ISGF3 activity, quantified from the EMSA (Fig. 2A), which was used as an input. The model simulations were fit to the median mRNA abundances for genes in each cluster at each time point for both stimuli (Fig. 2B) using a simplex search parameter optimization with 50 random initializations. To ensure robustness, the optimization workflow was conducted five times and the mean and associated standard error of the top 10 parameter sets from the five independent runs is reported. As expected, we were able to identify a parameter set that produced good fits for cluster 2 for both stimuli (RMSD = 0.119 ± 0.0001) (Fig. 2C). Then using the time course data of cluster 1, we identified a different parameter set that produced model simulations with a good fit (RMSD = 0.152 ± 0.0002). This suggests that the IFN-specific temporal dynamics of ISGF3 activity may be sufficient to mediate not only IFN-specific temporal control of ISG expression (cluster 2), but also activation of an IFN-β-specific gene expression program (cluster 1), without the need to invoke other pathways or transcription factors.

The parameter sets for the models for cluster 1 and 2 differed by 1.2x in the activation constant, by 1.5x in the degradation rate constant, and by 2x in the Hill coefficient. Using a sensitivity analysis where one parameter was changed by a multiplier, we found that for the cluster 1 model a four-fold increase or decrease in the activation constant, K_A,_ diminished the performance of the model only minimally (RMSD = 0.160 and 0.153, respectively) (Fig 2D). In contrast, a much smaller range (≥ 0.7x and ≤ 1.4x) for the values of the fitted Hill coefficient, n, was found to be necessary in both parameter sets. The cost function more than doubled (RMSD = 0.342) when decreasing the Hill coefficient in the cluster 1 model to 1.0, and more than tripled (RMSD = 0.380) when increasing it to 2.0 in the cluster 2 model. The mRNA degradation constant showed an intermediate sensitivity: a 0.7- and 1.4-fold change for model 1 and 2, respectively, resulting in a slightly increased RMSD (0.173 and 0.153, respectively). Taken together, this suggests the decoding of ISGF3 temporal dynamics relies on specific values of the Hill coefficient and, to a smaller extent, the mRNA degradation constant.

Given that the cluster 2 model fit poorly to cluster 1 data (RMSD = 0.432), we asked which parameter difference was critically important to distinguish cluster 1 from cluster 2 (Fig. 2E). We found that substituting the Hill coefficient alone was sufficient to improve the model performance to more closely simulate the cluster 1 data (RMSD = 0.183). Substituting both the Hill coefficient and degradation rate further improved the fit (RMSD = 0.152). The dose-response relationship of mRNA abundance against ISGF3 activity for the cluster 2 model is close to linear for the experimentally determined range of ISGF3 activity (Fig. 2F). For the experimentally measured maximum ISGF3 activity induced by IFN-λ3 (1.4 A.U., red dotted vertical line), the mRNA abundance is 62% compared to the maximum mRNA abundance seen with the measured IFN-β induced activity (2.5 A.U., blue dotted vertical line). This is in contrast to the cluster 1 model whose non-linear dose response results in only 31% mRNA abundance for 1.4 A.U. ISGF3 activity.

The modeling suggested an ultra-sensitive dose response curve for cluster 1 genes versus a linear dose-response curve for cluster 2 genes. Such non-linear dose responses may arise from multiple transcription factors binding and synergizing in the activation of the promoter [39, 40], or from chromatin looping to connect distal enhancers with promoters [41-43]. We investigated each possibility by examining STAT1 ChIP-seq data and identified STAT1 binding motifs within the measured STAT1 peaks. This analysis did not reveal an over-representation of multiple motifs in STAT1 peaks associated with cluster 1 genes (Fig. 2G). However, many more cluster 1 genes showed larger distances between STAT1 peaks and transcription start sites (TSSs), while for cluster 2 genes STAT1 peaks tended to be promoter-proximal (Fig. 2H). This analysis suggests the possibility that the ultra-sensitive dose-response behavior that is driving the type I-specific expression of cluster 1 genes may be mediated by the need to engage in chromatin looping events that connect distal ISGF3 binding to the TSSs of target genes.

### A model of the IFN-ISGF3 signaling network guides experimental studies

Since the IFN-type-specific temporal dynamics of ISGF3 appear to control IFN-specific gene expression levels, we examined the mechanisms for regulating ISGF3 dynamics. We leveraged an ODE model of the JAK-STAT signaling pathway that recapitulates IFN-α induced signaling in hepatocytes [37], which contains 41 molecular species connecting ligand-receptor interactions with nuclear ISGF3 activity (Supplemental Table 1). To adapt the model to IFN-β-induced ISGF3 signaling in epithelial cells we produced experimental data from time course measurements of the pathway regulators STAT1, STAT2, IRF9, SOCS1, SOCS3, and USP18 during IFN-β stimulation using immunoblot, EMSA, and qPCR assays (Fig. 3A). We sought to make the least number of parameter changes and thus implemented an iterative algorithm starting with only three systematically selected parameters to produce an optimal fit (Fig. 3B). However, key features were not captured, such as the potentiated phase of some of the nuclear active species (ISGF3 and pSTAT1), the late phases of the nuclear total species (STAT1, STAT2, IRF9), as well as the dynamics of most of the mRNA species (total RMSE = 73.26) (Fig. 3C). To improve the model fits, three additional parameters were systematically selected for optimization. We found that allowing 12 parameters to be estimated was sufficient to capture the potentiated phase of the nuclear active species and the late phase of the nuclear inactive species (total RMSE=43.72), but the 21-parameter estimation further improved the overall fit, particularly for STAT1 mRNA (total RMSE=35.18). The parameter changes (Supplemental Table 2) suggest that MLE-12 lung epithelial cells have altered nucleo-cytoplasmic transport to increase the nuclear residence time of ISGF3 and lower STAT1 and 2 expression levels than in the original model that was parameterized to hepatocytes. With the new MLE-12 adapted model we could now explore the circuit mechanisms that are key to IFN-β-specific ISGF3 dynamics.

**Figure 3.**
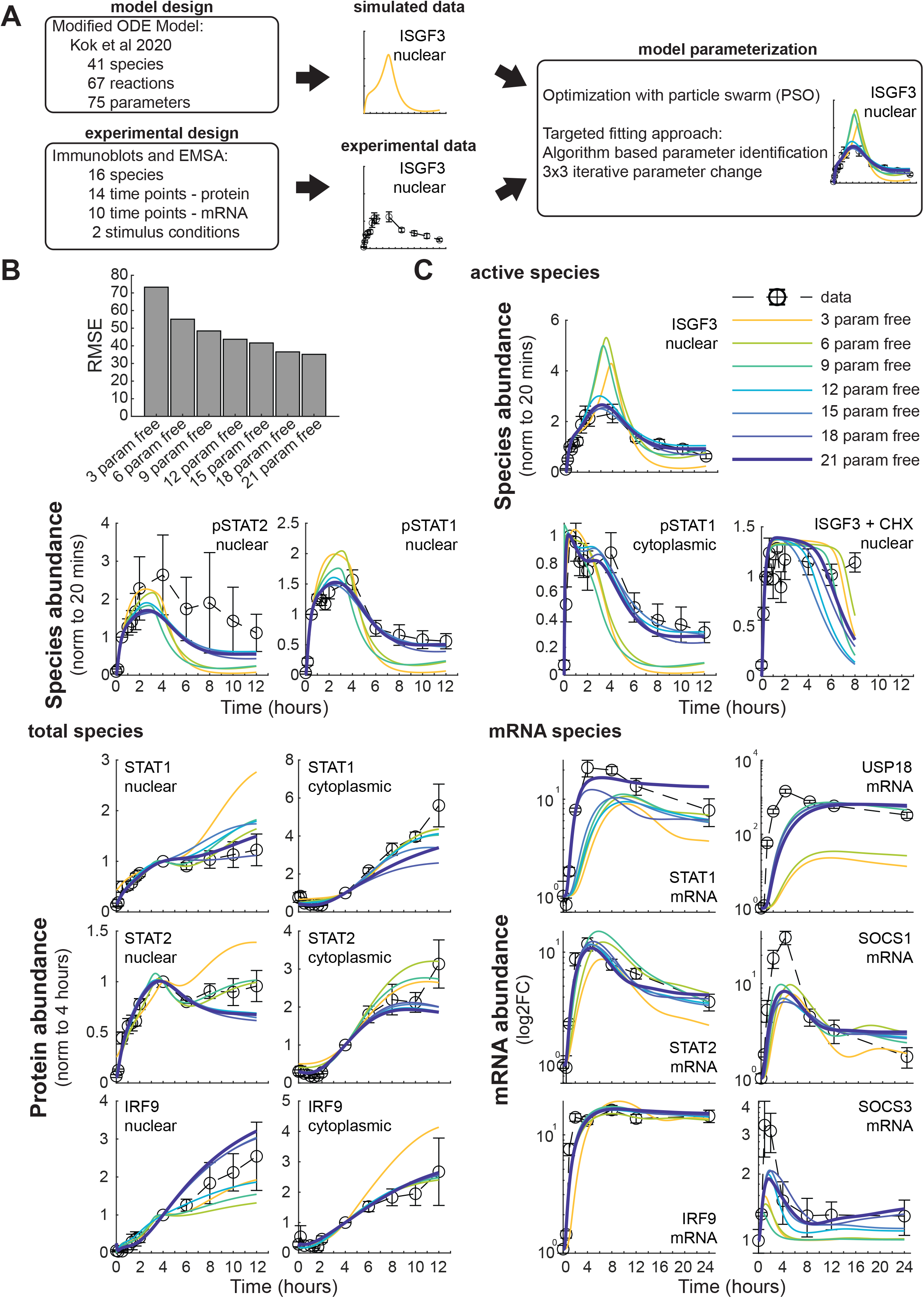
Applying a dynamical network model to IFN-β-induced lung epithelial cell signaling. A. IFN signaling network model design and parameterization. A previously published IFNα signaling model was modified and parameterized using a particle swarm optimization (PSO) to fit simulations to time course data of model species. To minimize alterations to the original model an iterative approach based on a 3-by-3 parameter identification and fitting was implemented. B. Bar graph of the root mean square error (RMSE) of optimally fit models compared to experimental data. Models, in which 18 or 21 of the 75 parameters are fit, provide an optimal fit. C. Comparisons of model simulations to time course datasets. Using quantified immunoblot and EMSA time course data (open circles with dotted black line) of nuclear and cytoplasmic active species (ISGF3, phosphorylated STAT1 (pSTAT1), phosphorylated STAT2 (pSTAT2)), total species (STAT1, STAT2, IRF9), and mRNA species (STAT1, STAT2, IRF9, USP18, SOCS1, SOCS3) during 10 U/ml IFN-β stimulation the best fit model (bold dark blue line) was determined. Visual inspection confirms that 18 or 21 parameter fitting results in satisfactory fits to the experimental data.

Using the signaling model adapted to IFN-β-responding MLE-12 lung epithelial cells, we explored how varying the dose of IFN-β stimulation would affect the temporal dynamics, and specifically, the potentiated phase of ISGF3 activity. First, we decreased the dose of IFN-β by two-, four-, ten-, or twenty-fold (Fig. 4A). While 1x IFN-β (50,000 molecules/cell) induced an initial activation phase up to one hour followed by a potentiated phase up to four hours, our simulations predicted a dose-dependency in the magnitude of the ISGF3 response. We plotted the ISGF3 activity during the initial activation phase at one hour and during the potentiated phase at four hours and found that the potentiated phase is reduced more drastically than the initial activation phase (Fig. 4B). For example, the model predicted that a ten-fold lower IFN-β dose would reduce the four-hour activity by about 2.3x, whereas the one-hour activity remained within 1.2x of the original, resulting in close to equal magnitudes at both timepoints. Experimentally, we tested three doses (10 U/ml, 1 U/ml, and 0.1 U/ml) in an ISGF3 time course experiment (Fig. 4C). Similar to the model predictions, ISGF3 activity trajectories were diminished at lower doses. The quantified experimental data showed a much more severe reduction in the four-hour potentiated phase than the one-hour activation phase such that both time points had close to equal ISGF3 activity at lower doses (Fig. 4D). This suggests that the IFN-β-specific potentiated phase of ISGF3 is dose-dependent and requires a higher dose-range than the initial activation phase which occurs also at lower doses.

**Figure 4.**
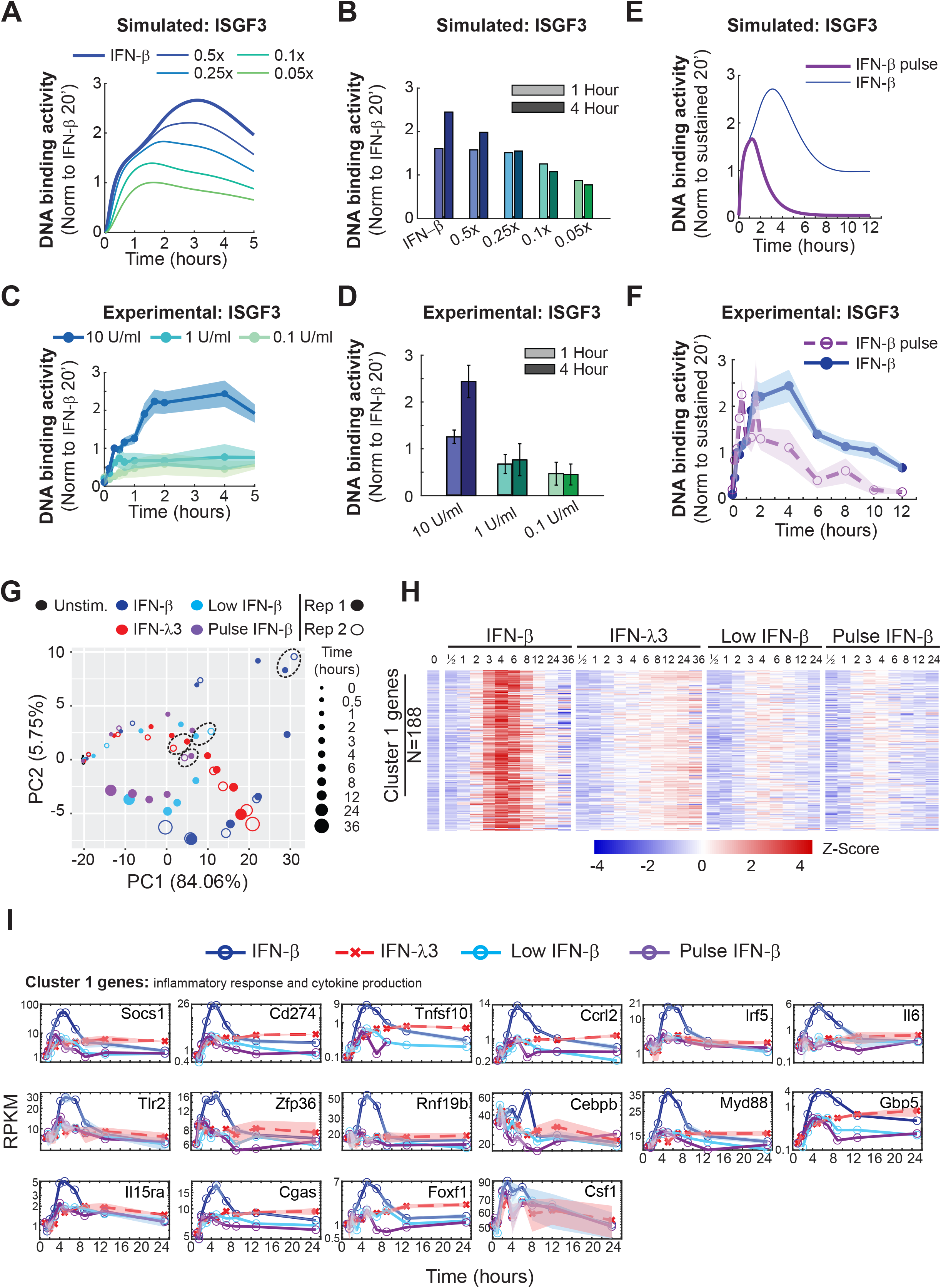
ISGF3 dynamics, which depend on the dose and duration of IFN-β stimulation, control expression of inflammation genes. A. Line graphs of model simulations of dose-dependent ISGF3 temporal dynamics. Using the adapted IFN-signaling model, simulations predict ISGF3 DNA binding activity at various IFN-β doses (thin solid lines) that are lower than the simulation that was fit to experimental data (thick blue line). B. Bar graphs of ISGF3 activity during the primary activation (one hour) and secondary potentiated (four hours) phases, as simulated by the model with indicated doses of IFN-β (panel A). C. Line graphs of experimental dose-dependent ISGF3 temporal dynamics. ISGF3 activity during 10 U/ml IFN-β stimulation was revealed by an EMSA and quantified. The SEM is depicted from at least two independent experiments. D. Bar graphs of ISGF3 activity during the primary activation (one hour) and secondary potentiated (four hours) phases, as experimentally determined for indicated doses of IFN-β (panel C). E. Line graphs of simulated ISGF3 activity during sustained (blue line) and a 15-minute pulse (purple line) of IFN-β. F. Line graphs of experimental ISGF3 activity during sustained (blue line) compared to a 15-minute pulse (dashed purple line) of 10 U/ml IFN-β. Data are from at least three independent experiments. G. PCA of IFN-induced transcriptomes with sustained high (10 U/ml, blue), low (1 U/ml, light blue), or pulse (15 minute of 10 U/ml, purple) IFN-β and IFN-λ3 (100 ng/ml, red) stimulus conditions. Trajectories of gene expression levels over time (increasing circle size) are depicted for two independent experiments (open and closed circles). The four-hour time point for each condition is indicated by dashed circles. H. Heatmap depicting the time course of scaled mRNA abundances of individual genes during IFN stimulus conditions eliciting stimulus-specific ISGF3 temporal dynamics. Mean expression levels for each gene in cluster 1 during IFN-β (10 U/ml), IFN-λ3 (100 ng/ml), low IFN-β (1 U/ml), or pulse IFN-β (15 minutes of 10 U/ml) are depicted in white while values above or below the mean are depicted in shades of red or blue, respectively. I. Line graphs of mRNA abundance time course induced by IFN-β, IFN-λ3, low IFN-β, or pulse IFN-β. Genes selected based on analysis of gene cluster-specificity and biological importance from Figure 1F are depicted here with connecting lines.

To determine if the ISGF3 potentiated phase is dependent not only on the dose but also the duration of stimulation, we compared model simulations of the sustained 1x IFN-β condition and a pulse stimulation where IFN-β was set to zero after 15 minutes (Fig. 4E). While the activation phase was similar in both conditions, the potentiated phase did not occur with the shorter stimulation time. Instead, ISGF3 activity rapidly decreased to baseline within six hours. We tested this prediction experimentally in MLE-12 lung epithelial cells stimulated with 10 U/ml IFN-β for 15 minutes (Fig. 4F). Similar to the model predictions, the experimental ISGF3 activity did not increase after the activation phase but diminished to basal activity by six hours. Together, the modeling and experimental results suggest that the IFN-β-specific potentiated phase of ISGF3, requires higher doses and longer durations of ligand exposure.

The above-described stimulation conditions allowed us to assess the functional consequence of the ISGF3 potentiated phase for gene expression. We isolated RNA from MLE-12 lung epithelial cells incubated with either low IFN-β (1 U/ml) or pulse IFN-β (15 minutes of 10 U/ml) for various durations and assessed the expression of the previously identified 345 IFN-β inducible genes by RNA-seq. Principal component analysis revealed that the response profile for neither condition converged with the response profile for sustained IFN-β after one hour, but clustered near the IFN-λ3-induced profile until four hours (Fig. 4G, dashed circles). After four hours, the pulse and low IFN-β-induced transcriptomes converge towards the unstimulated condition. When examining the previously identified IFN-β-specific gene expression program in cluster 1, the lower dose and shorter duration of IFN-β was not sufficient to induce these genes (Fig. 4H). Line graphs of cluster 1-specific inflammatory & adaptive immune response and cytokine production genes (e.g. Cd274, Il6, TNFSF10, CCRL2, Tlr2, Myd88, and IRF5) confirmed that only the sustained and high dose of IFN-β was able to induce them (Fig. 4I). This demonstrates that the ISGF3 potentiated phase induced with high and sustained IFN-β stimulation is needed to activate expression of specific genes involved in the regulation of inflammatory and adaptive immune responses, and cytokine production.

### A stimulus-contingent positive feedback loop interprets stimulus dose and duration

Since the ISGF3 potentiated phase has biological importance in regulating a subset of inflammatory genes, we investigated the mechanisms controlling this phase of the ISGF3 response. Given the timing of the response and the dependency on dose and duration, we hypothesized that a stimulus-contingent positive feedback loop was regulating the ISGF3-potentiated phase. To test this, the protein synthesis inhibitor cycloheximide (CHX) was incubated with MLE-12 lung epithelial cells stimulated with IFN-β to block induced feedback loops (Fig. 5A). Time course measurements of ISGF3 activity revealed that while the initial phase of ISGF3 activity was not affected by the CHX treatment, the potentiated phase, which starts after the first hour, was blocked. The level of ISGF3 activity induced during the activation phase was sustained throughout the remainder of the time course, suggesting that a positive feedback loop is required to amplify the ISGF3 response. To identify potential gene candidates for the positive feedback amplifier, we measured STAT1, STAT2, and IRF9 mRNA abundances, known positive regulators of IFN signaling, during an IFN-β stimulated time course using RT-qPCR (Fig. 5B). By 60 minutes, IRF9 mRNA had already increased about 8-fold, while STAT1 and STAT2 mRNA had doubled at that time. Measuring the nuclear STAT1, STAT2, and IRF9 protein, we also observed that IRF9 had the largest increase (7-fold) during the potentiated phase (100 minutes) with respect to the activation phase (30 minutes) compared to STAT1 (40% increase) and STAT2 (30% increase) (Fig. 5C). This suggests that IRF9 induction, and to a lesser extent STAT1 and STAT2 induction, acts as a critical positive autoregulation circuit for ISGF3 and is needed for the stimulus contingent positive feedback loop.

**Figure 5.**
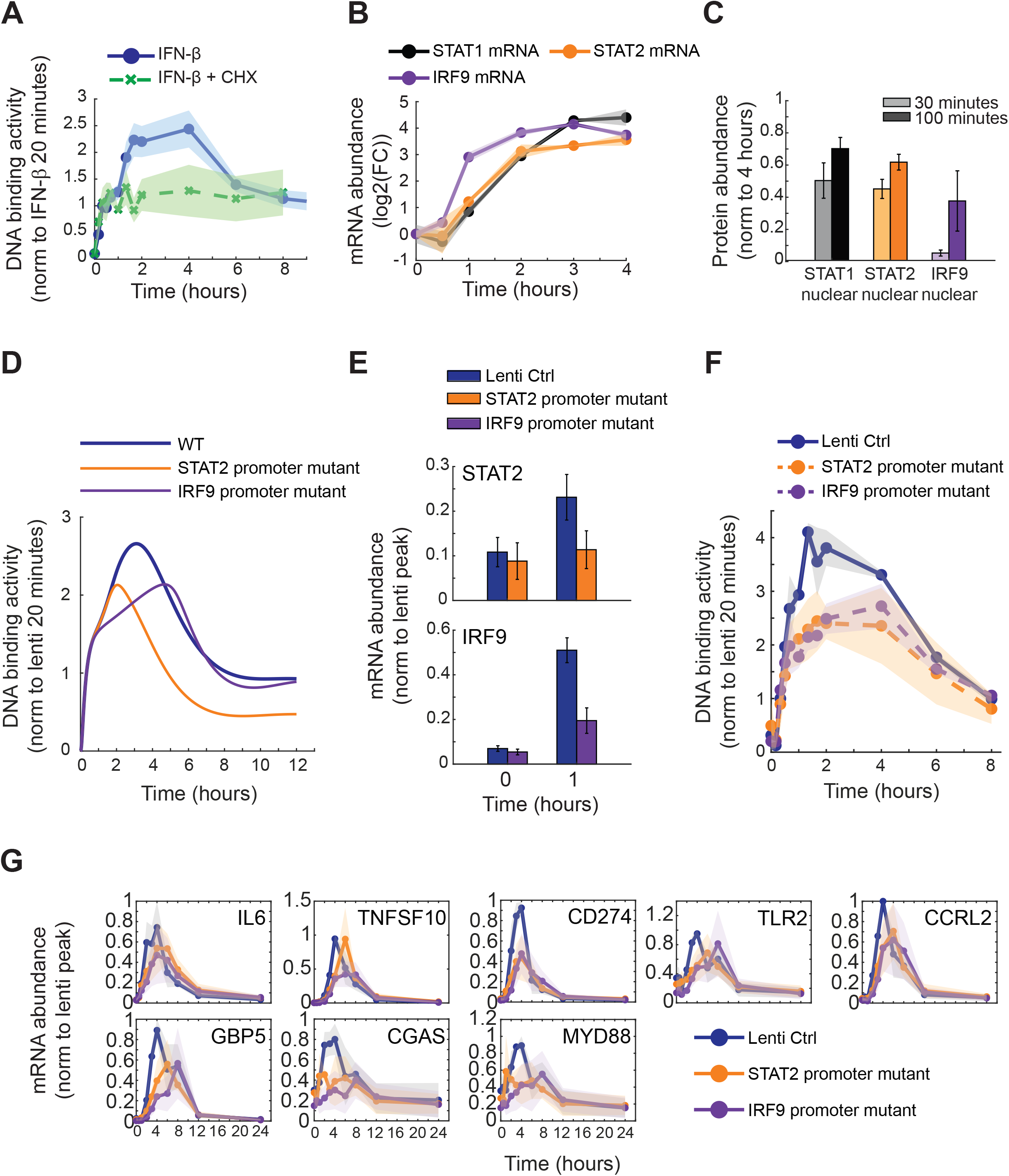
A stimulus-contingent positive feedback loop regulates the ISGF3 potentiated phase and is needed for ISGF3 potentiated-dependent gene induction. A. Line graphs of ISGF3 temporal dynamics during inhibition of protein synthesis. The temporal dynamics of ISGF3 activity when stimulated with 10 U/ml IFN-β with (blue line) or without (green dotted line) 10 μg/ml cycloheximide (CHX) was measured by EMSA. The SEM is depicted from at least three independent experiments. B. Line graphs of mRNA expression levels of positive regulators of IFN signaling. The temporal dynamics of STAT1 (black line), STAT2 (orange line), and IRF9 (purple line) mRNA induction during IFN-β stimulation (10 U/ml) were determined with a time course of gene expression levels measured using qRT-PCR. The SEM is depicted from at least four independent experiments. C. Bar graphs of nuclear STAT1, STAT2, and IRF9 protein levels during the primary activation phase (30 minutes) and start of the secondary potentiated (100 minutes) phase. Nuclear STAT1 (black bars), STAT2 (orange bars), and IRF9 (purple bars) protein abundances were measured using quantified immunoblots of MLE-12 lung epithelial cells stimulated with 10 U/ml IFN-β. Data was normalized to four hours when the protein levels are more abundant. The SEM is depicted from at least four independent experiments. D. Line graphs of model simulations of STAT2- and IRF9-feedback loop dependent ISGF3 temporal dynamics. Using the adapted IFN-signaling model (Figure 3), simulations predict ISGF3 binding activity when positive feedback loops for STAT2 (yellow line) and IRF9 (purple line) are inhibited compared to the simulation of the WT model (thick blue line). E. Bar graphs of mRNA gene expression levels of STAT2 and IRF9 in STAT2 and IRF9 promoter mutants. STAT2 and IRF9 mRNA abundances were measured using RT-qPCR of lentiviral control (blue bars), STAT2 promoter mutant (orange bars), or IRF9 promoter mutant (purple bars) MLE-12 lung epithelial cells stimulated with 10 U/ml IFN-β. Data was normalized to the peak expression level at 4 hours of the lentiviral control. The SEM is depicted from at least four independent experiments. Statistics were generated using a Student’s t-test. F. Line graphs of experimental STAT2- and IRF9-feedback loop dependent ISGF3 temporal dynamics. The temporal dynamics of ISGF3 activity in MLE-12 lung epithelial cells with mutations in the STAT2 promoter (orange dashed line), IRF9 promoter (purple dashed line), or a lentiviral control (black line) when stimulated with 2.77 U/ml IFN-β was measured with an EMSA. The SEM is depicted from at least two independent experiments. G. Line graphs of mRNA gene expression levels of important inflammatory genes in STAT2 and IRF9 promoter mutants. mRNA abundances of genes selected based on biological importance from Figure 1F is depicted from lentiviral control (blue bars), STAT2 promoter mutant (orange bars), or IRF9 promoter mutant (purple bars) MLE-12 cells stimulated with 10 U/ml IFN-β. Data was normalized to the peak expression level of the lentiviral control. The SEM is depicted from at least three independent experiments.

Using the MLE-12 IFN signaling model, we tested how IRF9 as well as STAT2 positive autoregulation during IFN-β stimulation affects ISGF3 activity (Fig. 5D). Model simulations predicted that when IRF9 induction is blocked the potentiated phase of the ISGF3 response is reduced, while the activation and late phase are similar to the control. Blocking the STAT2 positive autoregulation reduced the potentiated phase but also the late phase. In order to test the model predictions, the ISGF3 binding site within the IRF9 and STAT2 promoters in the MLE-12 lung epithelial cells were mutated using CRISPR-Cas9 gene editing. This significantly diminished IRF9 and STAT2 inducibility by IFN-β, as assessed by RT-qPCR (Fig. 5E). While the IRF9 promoter mutant and lentiviral control cells had similar basal levels of IRF9, at one hour of stimulation the abundance of IRF9 was reduced in the mutant cells compared to the control (p-value < 0.01). A similar reduction of STAT2 mRNA expression levels was detected in the STAT2 promoter mutant by one hour of stimulation (p-value < 0.05), suggesting diminished induction of the positive autoregulation in the IRF9 and STAT2 promoter mutants.

To examine the functional consequences of diminished autoregulation, the IRF9 and STAT2 promoter mutant cells were stimulated with IFN-β, and ISGF3 activity at various time points measured using EMSA (Fig. 5F). Similar to the model simulation, the potentiated phase was found to be diminished in both the IRF9 and STAT2 promoter mutants compared to the lentiviral control cells. To determine gene expression consequences, the mRNA of inflammatory instigator genes (e.g. IL6, TNFSF10) as well as inflammatory regulator genes (e.g. CD274, TLR2, CCRL2, GBP5, CGAS, and MYD88) from cluster 1 during IFN-β stimulation was measured with a time course using qRT-PCR (Fig. 5G). For the IRF9 promoter mutant, the amount of mRNA abundance was decreased in at least one time point for TNFSF10, CD274, CCRL2, GBP5, and CGAS compared to the lentiviral control (p-value <0.05). For many of the genes including TLR2, MYD88, and GBP5, the peak expression level was diminished and delayed to eight hours. The STAT2 promoter mutant also had a reduction in mRNA abundance for CD274, GBP5, and CGAS (p-value < 0.05), with shifts in peak expression for many of the genes compared to the lentiviral control. Taken together, this suggests a role for IRF9 and STAT2 positive autoregulatory feedback loops in controlling inflammatory gene expression responses during IFN-β stimulation.

## DISCUSSION

Here we report that in lung epithelial cells type I IFN-β induces an anti-viral gene expression program similar to type III IFN-λ3, and also a gene expression program that includes the prominent inflammatory instigators IL6, TRAIL, and CCRL2. Using experimental and mathematical modeling, we show that the type I IFN-specific gene expression program is regulated by the temporal dynamics of ISGF3. By adapting a mathematical model of the IFN signaling pathway to epithelial cells, we predicted and then experimentally confirmed the dose- and duration-dependency of the ISGF3 temporal dynamics necessary for the type I IFN-induced inflammatory gene expression program. We also found that the ISGF3-induced expression of its components, STAT2 and IRF9, constitute positive feedback loops that are important for regulating ISGF3 temporal dynamics and the gene activation of inflammatory instigators.

Our study emphasizes that the IFN gene expression response is not monolithic but depends on the type, dose, and duration of stimulus exposure. Previous studies have alluded to this by observing that type I and III IFN-induced gene expression in hepatocytes show different kinetics [18, 44]. Using IFNAR and IFNλR overexpressing cells it was found that IFN-type-specific kinetic differences were not due to the IFN dose or receptor abundance [45], but the underlying mechanisms remained obscure. Here, we also identified a group of genes (whose biological functions are primarily in cell-intrinsic innate defenses) in epithelial cells that show differential kinetics in response to type I and III IFNs. We also identified a group of genes (whose biological functions include cell extrinsic inflammation-induced immune responses) that show substantially different magnitudes such that they may be described as being expressed stimulus-selectively, specific to high dose IFN-β exposure. Furthermore, we address the underlying mechanism. Our study suggests that the stimulus-specificity in gene expression responses may be accounted for by stimulus-specific dynamics of the JAK-STAT/IRF signaling network and need not involve additional pathways that were previously implicated, such as MAPK or NFκB [34-36, 46]. We provide evidence that type I and III IFN stimulation result in distinct temporal dynamics of ISGF3; while initial activation is similar, only IFN-β-induced ISGF3 shows a potentiated phase. The response selectivity of cell extrinsic inflammation-induced immune genes due to differences in ISGF3 temporal dynamics could be recapitulated with a mathematical model by postulating a degree of ultra-sensitivity in ISGF3-promoter interactions. The fact that these genes have a larger distance between their transcription start sites and STAT1 binding sites, and thus may require DNA looping during the activation process suggests a plausible explanation for ultrasensitive, non-linear dose responses [41-43]. Nevertheless, our analysis also revealed substantial heterogeneity within the gene expression response, such that there may be multiple regulatory mechanisms. For example, a coherent feed-forward loop with ISGF3 and ISGF3-induced IRF1 may play a role [47]. Indeed, a finer gene expression analysis and more IFN stimulation conditions will likely identify additional gene expression patterns, as ultimately every immune response gene is probably subject to unique regulatory control mechanisms [48, 49].

Our study addresses the molecular mechanisms by which IFN-β-induced IGSF3 temporal dynamics are encoded. Prior mechanistic studies of IFN subtype differences have focused on IFN-receptor affinity measurements: IFN-β has high affinity (IFNAR2: 0.2 nM, IFNAR1: 50 nM) [14, 16], while type III IFN-λ3 binding affinities for its cognate receptors are weaker (IL10Rβ: 850 nM, IFNλR1: 16 μM). This could play a role in the differential cellular responses [50]. However, kinetic properties of receptor trafficking, receptor turnover and recycling, ligand half-life, and autoregulation mediated by feedback loops may be just as relevant.

Our iterative systems biology analysis using a quantitative dynamical systems model enabled our studies of ISGF3 regulatory mechanisms [37] and emphasizes the importance of the feedback loops in tailoring the gene expression response to type, dose and duration of exposure. The critical role of IRF9 in determining the magnitude of ISGF3 was previously described [38], but here we provide evidence that autoregulation of ISGF3 mediated by IRF9 and STAT2 results in a positive feedback loop that shapes the stimulus-specific dynamics of ISGF3. Positive feedback loops due to positive autoregulation in transcriptional network motifs are known to determine response times, signal sensitivity and amplification, and system bi-stability [51-53]. We demonstrate that autoregulation of ISGF3 results in positive feedback loop characteristics but also functions as a filter to distinguish stimulus duration and dose, akin to a coherent feedforward loop [54]. This occurs because active ISGF3 activates the expression of ISGF3 components (inactive ISGF3) that then require continued kinase activity to produce more active ISGF3 to induce the potentiated phase. A stimulus-contingent positive feedback loop governs in principle other immune response transcription factors, such as NFκB, which induces the expression of its components RelA, cRel and p50, as well as AP1. This induces the expression of its components Fos and Jun. Our study here may prompt an examination of their dynamical properties to determine whether they also function as filters of dose and duration potentially enabling the stimulus-specific activation of target genes. Finally, other feedback loops may also play a role in specifying the dynamic control of ISGF3, particularly at later times beyond the potentiated phase described here.

Our work characterizes a role for type I IFN-β in cell extrinsic inflammation through induction of inflammatory instigator genes in epithelial cells. We report that the gene expression of inflammatory regulators IL-6, TNFSF10, the gene that encodes for TNF-Related Apoptosis Inducing Ligand (TRAIL), and CCRL2 is tightly controlled and dependent on potentiated activity of ISGF3. Previous reports have described a link between type I IFNs and IL-6 and TNFSF10 in respiratory illnesses [55, 56], as the latter induces epithelial cell apoptosis [19, 57, 58]. CCRL2 is a non-signaling transmembrane receptor that presents chemerin, which recruits leukocytes. The detrimental effects of inflammation are prevented in CCRL2 deficient mice [59].

Precise control of IFN responses is critically important, as misregulation of IFNs can result in immunopathogenesis and autoimmunity or exacerbate the pathogenesis of respiratory viral infections [55, 56, 60-62]. While high IFN-λ1 levels were associated with lower SARS-CoV-2 viral loads and faster clearance in a cohort of COVID19 patients [55], a higher IFN-β-to IFN-λ1 ratio was linked to longer hospitalization. Similarly, in a SARS-CoV-1 infection model late activation of type I IFN contributed to lung pathology and morbidity [20]. This highlights the need to further explore type I IFN cell extrinsic roles during immunopathogenesis.

Misregulation of IFNs and their signaling responses leading to disease has also been linked to genetic variants. Inborn errors of immunity (IEI) have been identified for genetic variants of STAT1 and IRF9 with mutations causing either gain or loss of function [63]. Other pathway regulators have also been shown to have variants causing IEI such as JAK1, TYK2, IFNAR1, and IFNAR2 [63-65]. While IEI have been thought to be rare, common single nucleotide polymorphisms (SNPs) have also been found in TYK2 and IFNAR1 and are associated with improved protection from infection as well as increased risk of autoimmunity [66]. The prevalence of such SNPs may account for a wide range of IFN pathway responses among individuals [67, 68]. It will be of interest to delineate how such variants affect the engagement of the described positive feedback loop. Furthermore, the feedback expression of IRF9 and STAT2 may be subject to epigenetic regulation, such that exposure history could modulate pathway regulation and lead to highly individual immune responses. As the inflammatory gene cluster discriminates between modest and high levels of ISGF3 we would expect that there is more variation in the IFN-inflammatory response axis than the anti-viral defense functions of IFN.

Our work suggests that ISGF3 dynamic control may be key to understanding the plethora of specific IFN response functions. It suggests strategies to studying these further, leveraging mathematical modeling to develop increasingly reliable simulations that may be a tool for evaluating the potential impact of individual genetic and epigenetic differences for a new generation of precision medicine.

## MATERIALS & METHODS

### Cell Culture

MLE-12 lung epithelial cells were obtained from ATCC (#CRL-2110) and cultured in DMEM with 4.5 g/L glucose, L-glutamine, and sodium pyruvate (Corning #10-013-CV) supplemented with 10% fetal bovine serum, 100 IU/ml penicillin, 100 μg/ml streptomycin (Corning #30-002-CI), and 2 mM L-glutamine (Corning #25-005-CI) at 37°C at 5% CO_2_. For stimulations, MLE-12 cells were incubated with 1 U/ml or 10 U/ml recombinant mouse IFN-β (PBL Assay Science #12401-1) and 100 ng/ml recombinant mouse IL-28B/IFN-λ3 (R&D Systems #1789-ML-025/CF). For select experiments, cells were also incubated with 10 μg/ml cycloheximide (Sigma-Aldrich #C7698) or a vehicle (ethanol) control.

### Protein expression analysis by Immunoblotting

Protein from extracts was measured (BioRad DC™ Protein Assay Kit II #5000112) and equal amounts of protein were loaded into a 4-15% gradient polyacrylamide gel (BioRad TGX™ Precast Midi Protein Gel #5671085) and proteins separated based on molecular weight using electrophoresis at 110V. Proteins were transferred onto a PVDF membrane (Fisher Scientific Immobilon®-P IPVH00010) at 100V for one hour. The membrane was incubated for one hour in 5% bovine serum albumin (BSA) (Millipore Sigma #A3059) followed by overnight incubation at 4°C in a primary antibody solution. The following antibodies were used in this study: phospho-STAT1 (pY701.4A) mouse monoclonal IgG2aĸ targeting to phosphorylated Tyr701 (Santa Cruz Biotechnology #sc-136229), STAT1 p84/p91 (E-23) rabbit polyclonal IgG with epitope matching near the C-terminus (Santa Cruz Biotechnology #sc-346), STAT1 p84/p91 rabbit polyclonal IgG antibody (Cell Signaling #9172), phospho-STAT2 (Tyr689) rabbit polyclonal IgG antibody antibody with epitope matching at phosphorylated Tyr689 (EMD Millipore #07-224), STAT2 (D9J7L) rabbit monoclonal IgG antibody with epitope corresponding near Leu706 (Cell Signaling #72604), IRF9, clone 6F1-H5, mouse monoclonal IgG_2aĸ_ antibody with epitope matching the N-terminus (EMD Millipore #MABS1920), nuclear matrix protein p84 [EPR5662(2)] rabbit monoclonal antibody mapping with aa350-450 (abcam #ab131268), and αTubulin (B-7) mouse monoclonal IgG2aĸ antibody raised against epitope with aa149-448 (Santa Cruz Biotechnology #sc-5286). The membrane was washed in TBS-T (0.5% Tween-20) and incubated in a secondary antibody solution for one hour at room temperature. The following horseradish peroxidase (HRP) conjugated secondary antibodies were used for this study: mouse anti-rabbit IgG (L27A9) monoclonal antibody (Cell Signaling #5127), goat anti-rabbit IgG polyclonal antibody (Cell Signaling #7074S), horse anti-mouse IgG polyclonal antibody (Cell Signaling #7076), mouse anti-rabbit IgG monoclonal antibody (Santa Cruz Biotechnology #sc2357), bovine anti-mouse IgG polyclonal antibody (Santa Cruz Biotechnology #sc2380). The protein bands were resolved by chemiluminescence (Thermo Fisher Scientific SuperSignal West Pico and Femto Chemiluminescent Substrate #34080 and 34095).

### Electrophoretic Mobility Shift Assay

The protocol was used as previously described [69, 70]. In brief, nuclear fractions were collected by extraction using a Tris (250 mM Tris, 60 mM KCl, 1 mM EDTA) buffer solution. Equal amounts of nuclear protein were incubated for 30 minutes in a binding buffer (10 mM Tris-Cl, 50 mM NaCl, 10% glycerol, 1% NP-40, 1 mM EDTA, 0.1 mg/ml Poly(deoxyinosinic-deoxycytidylic) acid sodium salt) with 0.01 pmol P-32 radiolabeled oligo containing the ISRE consensus sequence (ISG15 loci; 5’-GATCCTCGGGAAAGGGAAACCTAAACTGAAGCC-3’; 5’- GGCTTCCAGTTTAGGTTTCCCTTTCCCGAGGATC-3’) or a NFY binding sequence (5’- GATTTTTTCCTGATTGGTTAAA-3’; 5’-ACTTTTAACCAATCAGGAAAAA-3’). Samples were loaded onto a 5% polyacrylamide gel (5% glycerol and TGE buffer [24.8 mM Tris base, 190 mM glycine, 1mM EDTA]). Bands were resolved using electrophoresis at 200V. Gels were dried and imaged using an Amersham Typhoon 5 laser scanner (GE Healthcare).

### Gene expression analysis by RT-qPCR

RNA was collected from cells lysed in TRIzol Reagent (Fisher Scientific #15-596-018) and isolated using the Direct-zol RNA Extraction Kit (Zymo Research #R2052). Equal amounts of RNA were reverse transcribed into cDNA using the Iscript™ Reverse Transcription Supermix (BioRad #1708841). Equal volumes of diluted (1:8) cDNA were loaded for qPCR analysis along with the SsoAdvanced Universal SYBR Green Supermix (BioRad #1725272) and 0.5 μM forward and reverse primers for target genes (listed below). Amplification curve thresholds were determined and selected for each primer set. mSTAT1 (5’-GGCCTCTCATTGTCACCGAA-3’; 5’-TGAATGTGATGGCCCCTTCC-3’); mSTAT2 (5’-GTAGAAACCCAGCCCTGCAT-3’; 5’-CTTGTTGCCCTTTCCTGCAC-3’); mIRF9 (5’-TCTTTGTTCAGCGCCTTTGC-3’; 5’-CTGCTCCATCTGCACTGTGA-3’); mSOCS1 (5’-CAACGGAACTGCTTCTTCGC-3’; 5’-AGCTCGAAAAGGCAGTCGAA-3’); mSOCS3 (5’-CTTTTCTTTGCCACCCACGG-3’; 5’-CCGTTGACAGTCTTCCGACA-3’); mUSP18 (5’-CTCACATGTTTGTTGGGTCACC-3’; 5’-TGAAATGCAGCAGACAAGGG-3’); mTNFSF10 (5’-GATGAAGCAGCTGCAGGACAAT-3’; 5’-CTGCAAGCAGGGTCTGTTCAAG-3’); mCD274 (5’-TCGCCTGCAGATAGTTCCCAAA-3’; 5’-GTAAACGCCCGTAGCAAGTGAC-3’); mCCRL2 (5’-GAGCAAGGACAGCCTCCGAT-3’; 5’-CCACTGTTGTCCAGGTAGTCGT-3’); mGBP5 (5’-TGCTGACATGAGCTTCTTCCCA-3’; 5’-TCATCGCTACCTTGCTTCAGCT-3’); mCGAS (5’-TGGGCACAAAAGTGAGGACCAA-3’; 5’-CGCCAGGTCTCTCCTTGAAAACT-3’); mMYD88 (5’-TCTCCAGGTGTCCAACAGAAGC-3’; 5’-TGCAAGGGTTGGTATAGTCGCA-3’); mTLR2 (5’-GGGGCTTCACTTCTCTGCTT-3’; 5’-AGCATCCTCTGAGATTTGACG-3’); mIL6 (5’-ACCAGAGGAAATTTTCAATAGGC-3’; 5’-TGATGCACTTGCAGAAAACA-3’); mGAPDH (5’-AGCTTGTCATCAACGGGAAG-3’; 5’-TTTGATGTTAGTGGGGTCTCG-3’).

### RNA Sequencing and Data Analysis

RNA was collected from cells lysed in TRIzol Reagent (Fisher Scientific #15-596-018) and isolated using the Direct-zol RNA Extraction Kit (Zymo Research #R2052). RNA libraries were generated using the KAPA Stranded mRNA-Sequencing Kit with KAPA mRNA Capture Beads (Roche #07962207001). Samples were enriched for poly(A)+ mRNA strands using mRNA Capture Beads. The captured mRNA was synthesized into complimentary DNA (cDNA). cDNA libraries were ligated using Illumina TruSeq single index adapters (Roche #KK8700), amplified, and quantified using a Qubit dsDNA Broad Range Assay Kit (Life Technologies #Q32850) and a Qubit 2.0 fluorometer. Sequencing was conducted on an Illumina HiSeq 2500 with single end 50 bp reads through the UCLA Broad Stem Cell Research Center. Sequencing reads were aligned and mapped onto the mm10 genome. The EdgeR package (Bioconductor 3.7) was used for differential gene expression analysis of raw read counts which were normalized using trimmed-mean of the mean (TMM) and filtered by the number of mapped reads and gene length (RPKM) of greater than 2. Normalized counts were applied to a negative binomial generalized linear model which was used to calculate Biological Coefficients of Variation. Pairwise dispersions between the normalized counts were calculated using the Cox-Reid Likelihood profile method. The differential expression testing was initiated using the likelihood ratio test (LRT), a form of ANOVA testing. Using the TREAT Method and Benjamini-Hochberg multiple test correction, significantly induced genes were selected based on differences in RPKM values at each time point were compared to values in unstimulated conditions using a false discovery rate (FDR) of less than 0.05 based on replicate data and fold-change greater than 2. PCA plots were generated using the prcomp function in R and plotted with ggplot2 [71]. Correlation plots were plotted using ggplot2. Heatmaps were generated using the pheatmap package.

### ChIP Sequencing and Data Analysis

The protocol was used as previously described [33]. In brief, cells were fixed with 2 mM disuccinimidyl glutarate for 30 mins followed by a 10-minute incubation in 1% formaldehyde. Cells are lysed (Buffer 1: 50 mM HEPES-KOH, 140 mM NaCl, 1mM EDTA, 10% glycerol, 0.5% NP-40, 0.25% Triton X-100, and 1X protease inhibitor cocktail (Thermo Fisher Scientific); Buffer 2: 10 mM Tris-HCl, 200 mM NaCl, 1 mM EDTA, 0.5 mM EGTA, and 1X protease inhibitor cocktail; Buffer 3: 10 mM Tris-HCl, 100 mM NaCl, 1 mM EDTA, 0.5 mM EGTA, 0.1% Na Deoxycholate, 0.5% N-lauroylsarcosine, and 1X protease inhibitor cocktail) and sonicated to shear the chromatin. The DNA-bound complexes were incubated with STAT1 p84/p91 rabbit polyclonal IgG antibody (Cell Signaling #9172) overnight at 4°C. STAT1 was immunoprecipitated by incubating for five hours at 4°C with Protein-G DynaBeads (Invitrogen #10004D). DNA bound to STAT1 complexes was extracted using 1% SDS and 0.6 mg/ml of Proteinase K (New England Biolabs #P81075) at 60°C overnight, followed by purification with Agencourt AMPure XP solid-phase reversible immobilization (SPRI) beads (Beckman Coulter #A63881). DNA was quantified using a Qubit dsDNA High Sensitivity Assay Kit (Life Technologies #Q32851) and a Qubit 2.0 fluorometer. Libraries were prepared using NEBNext Ultra II DNA Library Prep Kit NEBNext Multiplex Oligos (New England Biolabs) according to manufacturer instructions and sequenced with a length of 50bp on an Illumina HiSeq 2500 at the UCLA Broad Stem Cell Research Center. Sequencing data was aligned to mm10 genome as previously described [33]. MACS2 version 2.1.0 was used to identify peaks for each sample using pooled input samples as control with FDR < 0.01, and extension size of average fragment length [72]. STAT1 motifs were identified by searching the HOMER database for position weighted matrix files of STAT/GAS and ISRE/IRF motifs [73]. ISRE/IRF motifs included motifs for T1ISRE, ISRE, IRF2, IRF1, IRF3, and IRF8. STAT/GAS motifs included motifs for STAT1, STAT3+IL-21, STAT3, STAT4, and STAT5. A variant GAS motif in the promoter of Irf1 (GATTTCCCCGAATG) known to be bound by STAT1 was also included in the search as a GAS motif [74]. Peak to gene distances were calculated with bedtools, using the TSS of the longest isoform [75].

### Gene Ontology Analysis

The maximum RPKM value and gene names for the selected 188 genes in cluster 1 were analyzed using the Panther Classification System Database DOI: 10.5281/zenodo.4495804 Released 2021-02-01 [76]. The genes were mapped to the mm10 genome and analyzed using the statistical overrepresentation test which determines overrepresentation of gene ontology annotations in cluster 1 compared to all expressed genes using the Fisher’s Exact test and FDR correction. A list of GO biological process terms and IDs as well as associated genes was generated. Each GO term was categorized into 10 groups based on function. To determine if a function was enriched in a gene cluster the fraction of GO terms for each biological function compared to the total number of terms for the cluster was calculated and plotted as a pie graph. Similar analysis was conducted on the 157 genes in cluster 2.

### Lentiviral Cloning, Transfection, and Infection

The cloning strategy for the lentiCRISPRv2 plasmid was used [77, 78]. Briefly, the following oligos were annealed and cloned into a BsmBI (Thermo Fisher Scientific #FD0454)-digested lentiCRISPRv2 vector: STAT2 promoter mutant (5’-CACCGAGGGAAAGGAAACTGAAACC; 5’- AAACGGTTTCAGTTTCCTTTCCCTC-3’); IRF9 promoter mutant (5’- CACCGACTCAGACCACGTGGTTTCT; 5’-AAACAGAAACCACGTGGTCTGAGTC-3’).

Plasmids were transformed into One Shot™ Stbl3™ Chemically Competent bacteria (Thermo Fisher Scientific #C737303). Single colonies were selected and inoculated in LB Broth. Plasmids were isolated from expanded bacterial colonies using the ZymoPURE™ II Plasmid Kit (Zymo Research #D4203). Ten μg of isolated plasmid was transfected into HEK293T cells using Lipofectamine® 2000 Transfection Reagent (Invitrogen #11668-019). Two days following transfection, viral particles were isolated from the HEK293T conditioned media. MLE-12 lung epithelial cells were infected with the purified viral particles for 24 hours, followed by 2 μg/ml puromycin antibiotic selection.

### Mathematical Modeling of interferon-induced gene expression

An ordinary differential equation (ODE) mathematical model was built to simulate RNA abundance levels as a function of transcription factor binding and RNA degradation. Briefly, the median RPKM values for all genes within each cluster from the RNA sequencing data was calculated for each time point. The temporal gene expression trajectory of the median values was generated using a modified Akima interpolant fit (MATLAB). The model was composed of four parameters (e.g., k_act_, activation rate; k_deg_, mRNA degradation rate, K_a_, association constant, and n, Hill coefficient). The model was parameterized by model fitting to the gene expression trajectories normalized to max peak. For optimization, the simplex search method (e.g., fminsearch) was implemented using an objective function of the sum of RMSD values calculated at each time point. 50 multiple random seeds for the initial parameter values were used to optimize over a larger parameter space. The parameter optimization was run 5 times and the top 10 parameter sets with the lowest RMSD values were averaged and used as the optimized model parameter values. Heatmaps and line graphs were plotted using MATLAB.

### Mathematical Modeling of the interferon signaling network

We used the previously publish model of interferon signaling [37], with minor modifications. We replaced the equation for the active receptor complex formation by

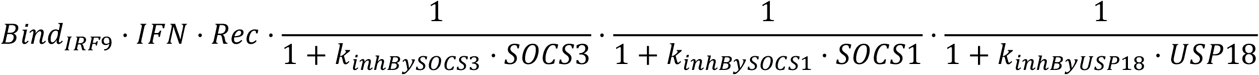

instead of

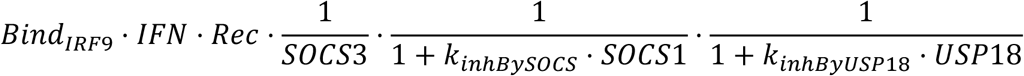

to be able to simulate the model in presence of cycloheximide which blocks protein synthesis, leading to SOCS protein decay, ultimately converging to 0. To better fit our data, we moved the delay term prior to mRNA synthesis instead prior to protein synthesis.

#### Parameterization

There are 41 molecular species in the mathematical model (Supplemental Table S1). The model parameters (Supplemental Table S2) were fit to the data, optimizing parameter value as well as which parameter to change iteratively 3 by 3 in order to adjust the minimal number of parameters allowing for a good fit.

##### Pseudocode

1. Initialize set of indices I_-1_ = {}
2. For k ranging from 0 to …
  a. Cost_k_ = ∞
  b. For run ranging from 1 to 20
    i. Optimize the cost function for θ = (i_3k+1_, i_3k+2_, i_3k+3_, v_1_, …, v_3k+3_) using particle swarm optimization with a number of particles equals to 10x the size of θ (= 10 x (3k+6)).
      - i_3k+1_, i_3k+2_, i_3k+3_ correspond to the indices of additional parameters to change,
      - v_1_, …, v_3k_ corresponds to the values for previously chosen parameters i.e. with indices from I_k-1_,
      - v_3k+1_, v_3k+2_, v_3k+3_ corresponds to the values of parameters i_3k+1_, i_3k+2_, i3k+3.
    ii. If cost_run_ < Cost_k_:
      - Ik = Ik-1 ∪ {i3k+1, i3k+2, i3k+3}
      - V_k_ = (v_1_, …, v_3k+3_)
      - Cost_k_ = cost_run_
  c. If Cost_k_ > Cost_k-1_
    i. Run step b. with more particles number: 20x size of θ.
3. Return I_k_ and V_k_

The cost function was the sum of the mean square error of each individual experiment, as follows:

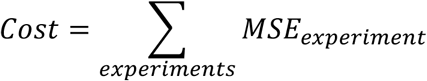

with

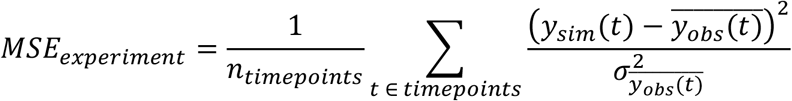

and with y_sim_ corresponding to simulated outputs for a specific parameter set.

#### Implementation

The modeling and parameter optimization were done in MATLAB R2018a using the particleswarm function implemented in MATLAB’s optimization toolbox.

## ACKNOWLEDGEMENTS

We thank Simon Mitchell for his discussions during the initial phase of the project.

## AUTHOR CONTRIBUTIONS

Catera L. Wilder: Conceptualization, Investigation; Project administration; Methodology; Formal analysis; Visualization; Writing – original draft

Diane Lefaudeux: Methodology; Formal analysis; Writing – review and editing

Raisa Mathenge: Formal analysis

Kensei Kishimoto: Investigation; Formal analysis; Writing – review and editing

Alma Zuniga Munoz: Investigation

Minh Nguyen: Formal analysis

Aaron Meyer: Supervision; Writing – review and editing

Quen J. Cheng: Investigation; Formal analysis; Writing – review and editing

Alexander Hoffmann: Conceptualization; Funding acquisition; Supervision; Writing – original draft

## SUPPLEMENTAL TABLES

Supplemental Table S1: Molecular species of the mathematical model.

Supplemental Table S2: Reactions and parameters of the mathematical model. Each row represents a molecular reaction. The parameters and values are indicated in each column for the original value and confidence interval (C.I.) from the (Kok et al 2020) IFNα signaling model as well as the estimated parameters for each model simulation.

**Supplemental Figure 1.**
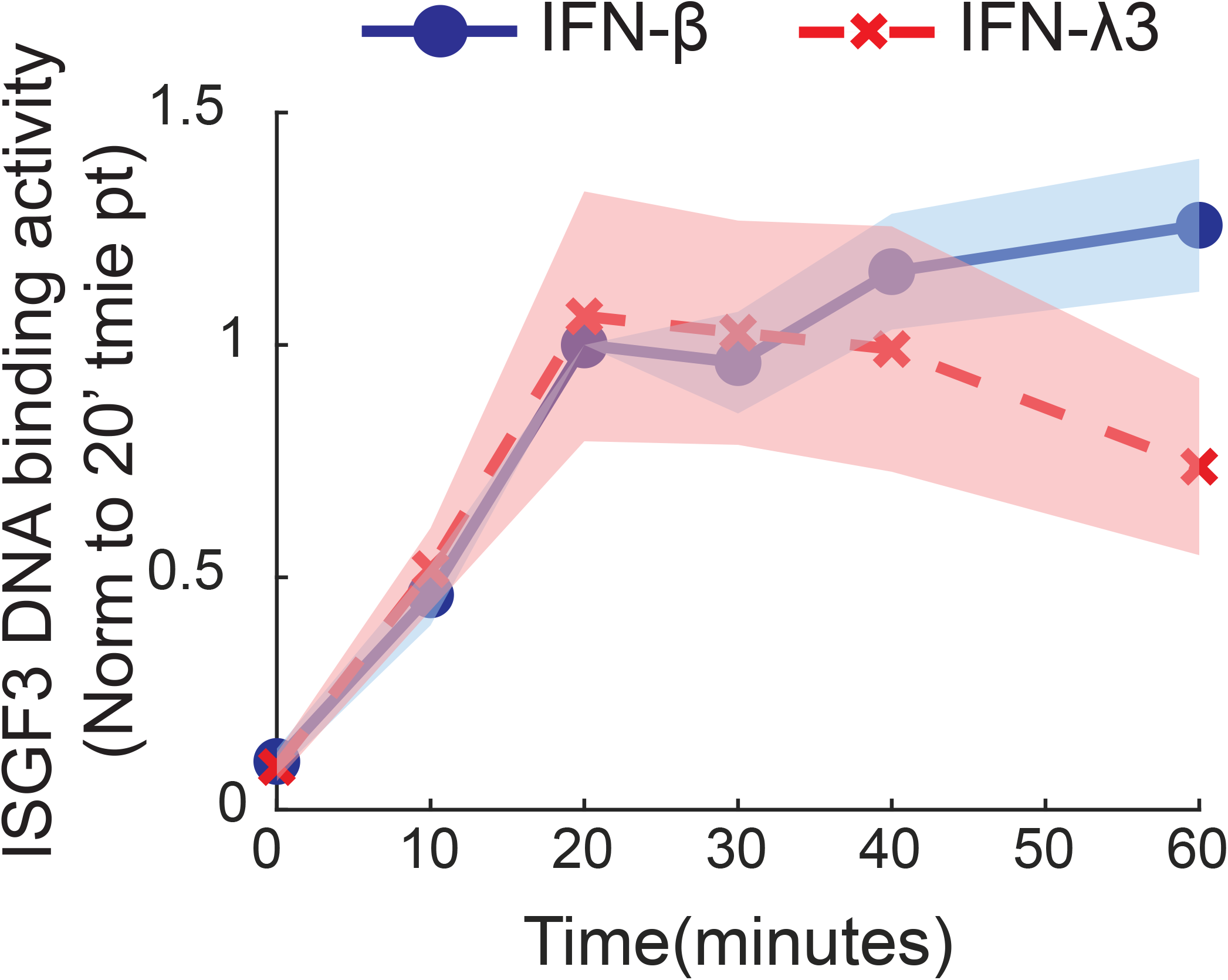

